# Damage of Major South American Lepidopteran Soybean Pests

**DOI:** 10.1101/2022.06.24.497488

**Authors:** Pablo Daniel Carpane, Matías Llebaria, Ana Flavia Nascimento, Lucía Vivan

**Affiliations:** Bayer CropScience, Fontezuela, Buenos Aires, Argentina; Universidade Federal de Uberlândia, Fundação Mato Grosso. Av. Antônio Teixeira dos Santos, 1559. Parque Universitário. Rondonópolis, MT, Brasil

## Abstract

Lepidopteran pests are major factors limiting soybean productivity in South America. In some cases, the control of these species requires the use of foliar insecticides. For a sustainable use of these insecticides, they should be sprayed when insect population sizes reach an economic threshold. Since this estimation requires to determine the consumption of different species, this work aimed to integrate all the main factors, studying the consumption of small-and medium-size larvae of major lepidopteran pests to vegetative and reproductive tissues on Bt and non-Bt soybeans. The damage to vegetative tissues was tested in detached-leaf assays in grow chambers, and to reproductive structures was measured in greenhouse with infestation at early (flowering) and mid reproductive (mid grain filling) stages. Based on the feeding behavior of the species tested, they were cast in four groups: a) *A. gemmatalis* and *C. includens*, defoliating only the RR variety with the lowest consumption of foliar area; b) *S. eridania*, defoliating both RR and IPRO varieties, consuming twice than the species mentioned above; c) *H. armigera*, defoliating and being the most damaging species to pods in the RR variety; d) *S. cosmioides* and *S. frugiperda*, defoliating and damaging pods in both varieties. The species differed in their ability to feed on IPRO varieties, so a different economic threshould could be considered. This clasification contributes to a recommendation of insecticide use sustainable, taking into account the behavior of these species that are major soybeans pests in South America.

## Introduction

Soybean is one of the most important row crops globally, with around 350 million t harvested per season as 2019, from which South American countries (mostly Brazil, Argentina, Paraguay, and Uruguay) contribute to around 50% of the global production (FAOSTAT 2021). Insect pests are one of the most important factors limiting soybean productivity in South America. Indeed, the lepidopteran species *Anticarsia gemmatalis, Chrysodeixis includens, Helicoverpa armigera, Spodoptera cosmioides, S. eridania* and *S. frugiperda* are considered important soybean pests based on their high prevalence and their impact on yield (Hoffmann-Campo et al. 2000, Santos et al. 2005, Czepak et al. 2013, Justiniano et al. 2014, Blanco et al. 2016), and can be found in field in complexes of more than one species of different relative size.

Control of these lepidopteran species relies mostly on two pillars: use of Bt soybeans and spraying of foliar insecticides. Intacta (Bt) soybeans are currently the main component to control damage of lepidopterans pests in soybean, providing protection from damage of *A. gemmatalis, C. includens* and *H. armigera* (MacRae et al. 2005, McPherson and MacRae 2009, Yu et al. 2013) and suppression against *Spodoptera* species (Bernardi et al. 2014, Bortolotto et al. 2014, Murúa et al. 2018), whose control might require insecticide sprays if their populations are large enough to cause economic damage. In addition, refuge areas may also need to be sprayed if insect pressure becomes high (IRAC 2021). However, foliar insecticides need to be sprayed in a environmentally and economically acceptable way, compatible with IPM principles (Forrester et al. 1993, McCaffery 1998, Mc Pherson and MacRae 2009). According to this principle, insecticides should be sprayed only when insect population sizes are equal to their economic threshold (ET), which considers the time taken to implement control measurements before the damage reaches the economic injury level (EIL), or damage severity that will cause economic loss to the crop (Pedigo et al. 1986).

The first step to estimate EIL is to characterize the consumption of different pest species (Hutchins et al. 1988). In this sense, some work has been done in the past with major soybean pests, which usually did not compared hosts or species (Reid and Greene 1973, Gamundi 1988, Barrionuevo et al. 2012, Favetti et al. 2015), studied the consumption of different host plants by the same insect species (Kidd and Orr 2001, Bavaresco et al. 2003, Barros et al. 2010, Montezano et al. 2014, 2019, Naseri et al. 2014, Moonga and Davis 2016, Specht and Roque Specht 2016, Gomes et al. 2017, Silva et al. 2017a), have focused on soybeans with a reduced range of insect species (Silva et al. 2017b, Specht et al. 2019), or compared Bt and non-Bt soybeans in a reduced number of species (Yu et al. 2013, Bernardi et al. 2014, Adams et al. 2016, Silva et al. 2016). The most complete characterizations that we are aware of focused on small larvae feeding vegetative tissues on non-Bt soybeans (Boldt et al. 1975, Bueno et al. 2011, 2013) or compared to Bt soybeans (MacRae et al. 2005, Murua et al. 2018). However, since most of the species mentioned above also consume soybean reproductive tissues (Fitt 1989, Hoffmann-Campo et al. 2000, Santos et al. 2005, Ramírez y Gómez 2010, Azambuja et al. 2015, Silva et al. 2017b), Intacta soybeans are widely prevalent in South America, and usually late-stages larvae move from weeds or neighbor crops (Azambuja et al. 2015) prompted us to integrate all these aspects. For this reason, this work aimed to characterize and compare the consumption of small-and medium-size larvae of *Anticarsia gemmatalis, Chrysodeixis includens, Helicoverpa armigera, Spodoptera cosmioides, S. eridania*, and *S. frugiperda* feeding on vegetative and reproductive tissues on Bt and non-Bt soybeans, for a more precise characterization of EIL on soybean crops.

## Material and Methods

### Insects and plants

Insect colonies of *Anticarsia gemmatalis, Chrysodeixis includens, Helicoverpa armigera, Spodoptera cosmioides, Spodoptera eridania* and *Spodoptera frugiperda* were obtained from field-collected insects from different areas of Brazil (collecting around 200 insects per location and species) on non-Bt soybean crops, and maintained at laboratory in controlled conditions (25°C, RH of 70%, photoperiod of 14:10 [L:D] h). Colonies incorporated new insects on a yearly basis to reduce inbreeding. *H. armigera* species status was confirmed according to Arneodo et al. (2015).

The Bt soybean cultivar M7739IPRO (containing the event MON87701, which expresses the Cry1Ac protein conferring control of some lepidopteran species at larval stages) and the nearly isogenic cultivar BMX Desafio RR (8473RSF) were used.

### Larval consumption at vegetative stages

Soybean leaflets were excised daily from plants at around V6-V8 (Fehr and Caviness, 1977) from the middle third of the plants, cleaned for 2 minutes in 5% Clorox, rinsed with water, and dried with paper towels before placing them in 10-cm diameter Petri dishes (one leaflet per dish). Two larvae of the desired growth stage were then placed in the Petri dishes using paint brushes. Larvae of L1 and L3 growth stages were tested. Water was provided by placing small wet cotton bolls in each Petri dish to avoid dehydration. This experiment was run in a growth chamber with the same settings for insect colony rearing.

After the infestation, the following activities took place on daily ion each Petri dish: determination of insect survival, measurement of foliar area consumed as percentage of total leaf area (Ortega et al. 2016), and replacement of leaflets if area consumed has exceeded 50%. This procedure continued until all larvae had pupated or died. In cases where no consumption was detected after a two-day period, leaflets were replaced anyway to avoid potential effects of dehydration upon foliar area consumption. Pupal weight (mg) was measured individually at the end of larval growth.

The percentage of defoliation was translated into area (square centimeters) by measuring the area of 20 representative leaflets per variety. These measurements were performed by taking pictures of leaflets placed beside a paper sheet containing a square of known area. The area of each leaflet was then measured in ImageJ (Abramoff et al. 2004) using the square for calibration.

### Larval consumption at reproductive stages

Plants were grown in 12-liter plastic pots in a greenhouse at the desired growth stage. Each pot was placed on a tray, and was irrigated by adding water to the trays, avoiding the potential effect of high moisture upon larval mortality. At either early (R2) or late (R5) (Fehr and Caviness, 1977) reproductive growth stages, plants were infested with two larvae of the desired growth stage (L1 or L3) using paint brushes. Each plant was covered individually in voile to avoid larval movement between plants.

Fifteen days after the infestation, foliar area consumed, number of damaged and damaged pods and grain yield (g) were measured per plant basis. The foliar area consumed was estimated as the difference between the average foliar area of uninfested plants (ten plants per soybean variety) and the remaining foliar area of each infested plant. Both areas were estimated by taking pictures of leaflets beside a square of known area as described above.

### Experiment design and statistical analyses

Each Petri dish (for vegetative stage) or pot (for reproductive stage) was considered a replication. The number of replications was 30 for vegetative stages and 15 for reproductive stages. Petri dishes and pots were arranged randomly in the growth chamber and greenhouse respectively. The total number of treatments (24) consisted of the interaction of three factors: six insect species, two soybean varieties (IPRO and RR), and two larval growth stages (L1 and L3).

Statistical analyses were performed using Infostat (Di Rienzo et al. 2018). Predicted values were compared using DGC test (Di Rienzo et al. 2002), with a significance level of 10% for all variables.

Insect survival was analyzed according to Kaplan-Meier for the whole set of treatments. In cases where the p value was below 0.10, the most divergent treatment was determined visually and excluded from the following analysis. This process was repeated until p value exceeded the value of 0.10, indicating that survival of the remaining treatments was similar.

The number of pupae and the proportion of damaged pods were analyzed using generalized and mixed linear models. A binary distribution (pupated or not) with a logit linkage function were used for the number of pupae, being insect species, larval size and soybean variety considered fixed effects. Results were expressed as the proportion of larvae reaching pupal stage out of the 30 larvae infesting soybean leaflets. A binomial distribution was used for the number of damaged pods, adjusting the proportion using the total number of pods per pot.

Foliar area consumed, pupal weight, proportion of damaged pods and grain weight were analyzed using generalized linear models, being insect species, larval size and soybean variety considered fixed effects. Foliar area consumed was also analyzed using a linear regression.

## Results

### Survival of Larvae

The survival of larvae (Figure 1) varied across species. In *A. gemmatalis* (HA), the initial analysis indicated differences among Stage x Variety combinations (p<0.0001), due to a lower survival in IPRO. When the analysis was repeated in each variety, there was no difference in survival between L1 and L3 in IPRO (p=0.5171) and in RR (p=0.2881). In *C. includens* (CI) the same initial pattern was seen between varieties (p<0.0001), but L3 survived more than L1 in both IPRO (p=0.0251) and RR (p=0.0003). In *H. armigera* (HA) the initial difference (p<0.0001) was due to soybean varieties, while in the split analysis there were no differences between stages in IPRO (p=0.6807), and weakly in RR (p=0.0781), in this case due to no mortality in L1.

**Figure 1:**
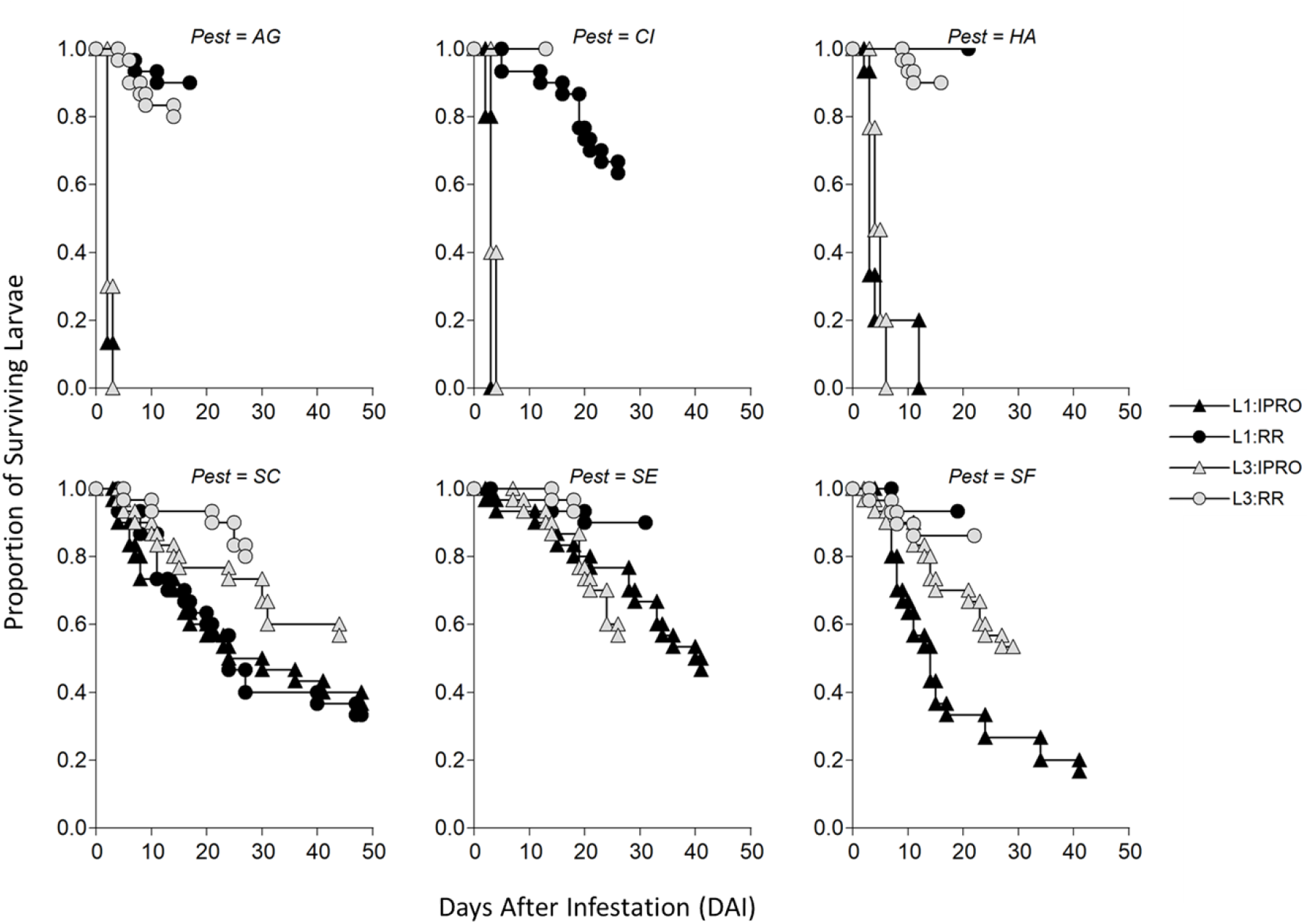
Proportion of surviving *Anticarsia gemmatalis* (AG), *Chrysodeixis includens* (CI), *Helicoverpa armigera* (HA), *Spodoptera cosmioides* (SC), *S. eridania* (SE), and *S. frugiperda* (SF) larvae over time (days) infesting leaf discs of IPRO or RR soybean varieties with larvae at L1 or L3 growth stages.

The pattern of survival in *Spodoptera* species was more complex than in the other species tested. In *S. cosmioides* (SC) the initial analysis showed significance (p=0.0013), being the main difference between larval stages, having L1 lower survival than L3. The analysis split by Stage indicated no differences in survival between varieties in L1 (p=0.3859), and a somewhat significant (p=0.0610) lower survival in IPRO by L3. In *S. eridania* (SE) the first analysis indicated a lower survival by larvae feeding IPRO. The split analysis per variety indicated no effect of larval stage in IPRO (p=0.6615) and RR (p=0.6382). Lastly, in *S. frugiperda* (SF) there was also a lower survival in larvae feeding in IPRO than in RR but, while there were no differences in survival in RR (p=0.3742), L3 survived more than L1 in IPRO (p=0.0037).

### Proportion of pupae

The proportion of larvae reaching pupal stage had a significant effect of Species (p=0.0018), and the interactions Species x Variety (p<0.0001) and a weak interaction (p=0.0708) of Species x Stage x Variety. The single effects Stage (p=0.9971) and Variety (p=0.9615), as well as the interactions Stage x Variety (p=0.9998) and Species x Stage (p=0.6192) were not significant. Figure 2 describes the triple interaction, showing that the Species effect and the Species x Variety interaction were because of no larvae of *H. armigera, A. gemmatalis* or *C. includens* reached pupal stage, while the species of *Spodoptera* did so. In this sense, all three *Spodoptera* species differed in this aspect: in *S. cosmioides* L3 feeding on RR had the highest proportion of pupae compared to the other ones, *S. eridania* survived more in RR than in IPRO without effect of larval size, and *S. frugiperda* had a higher proportion of pupae in RR than in IPRO, with L3 surviving more than L1 in this latter variety.

**Figure 2:**
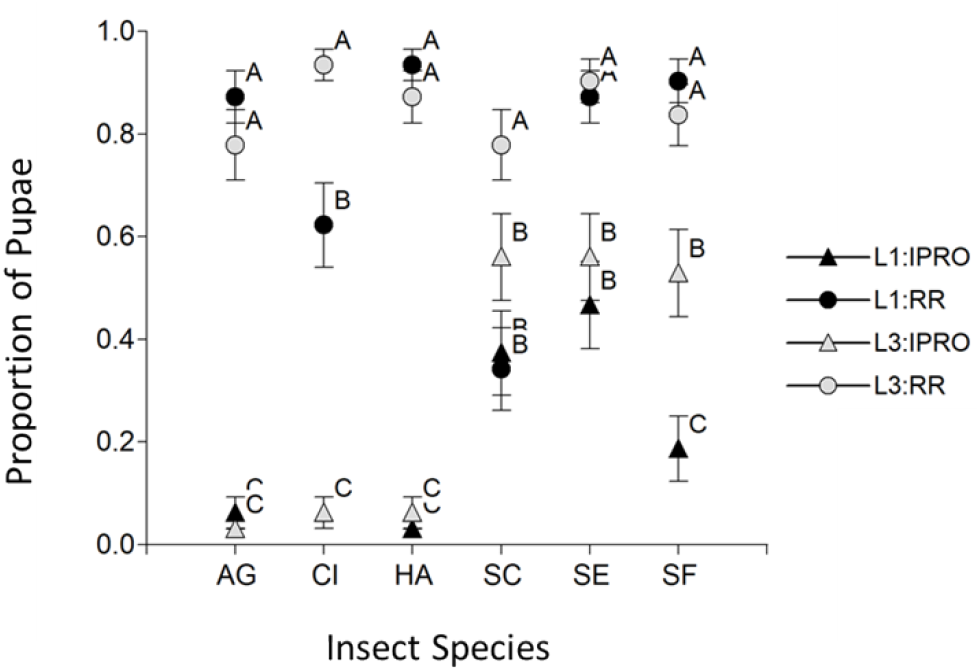
Proportion of *Anticarsia gemmatalis* (AG), *Chrysodeixis includens* (CI), *Helicoverpa armigera* (HA), *Spodoptera cosmioides* (SC), *S. eridania* (SE), and *S. frugiperda* (SF) larvae reaching pupal stage infesting leaf discs of IPRO or RR soybean varieties with larvae of L1 or L3 growth stages. Values with the same letter are not significantly different according to contrasts in the mixed model test (α = 0.10). Bars indicate standard error of the mean.

### Duration of Larval Period

Because no larvae of *A. gemmatalis, C. includens* and *H. armigera* having access to IPRO variety reached pupal stage (Figure 2), the duration of larval period was analyzed in two parts: a) These three species were analyzed to understand the effect of Stage and Species and their interaction, and b) The *Spodoptera* species were analyzed to see the effect of Stage, Species, Variety and their interaction. In the first comparison (Figure 3 Left), there was a significant effect of Species, Stage, and their interaction (p<0.0001 in all the cases). The Species effect was because of a longer larval period of *H. armigera*, and then by *C. includens*, and the interaction to the relatively longer length of larval period in L1 of *C. includens*. The analysis for *Spodoptera* species showed signifficant efects of Pest, Stage, Variety and Pest x Variety (p<0.0001 in all the cases) and Stage x Variety (p=0.0025) and for Pest x Stage x Variety (p=0.0006). The Variety effect was due to a longer duration of larval period in IPRO than in RR variety, and the triple interaction was because this delay differed across species. In *S. cosmioides* increase in duration of larval period was seen only on L3, in *S. eridania* in the same amount in both larval sizes, and in *S. frugiperda* mostly in L1.

**Figure 3:**
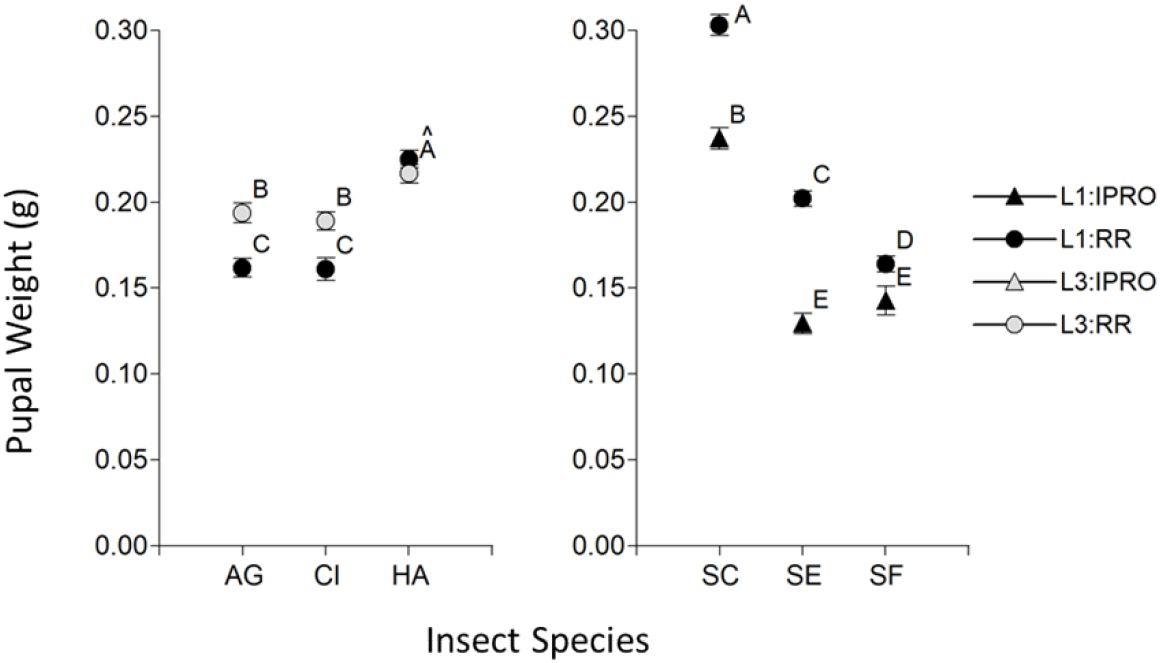
Duration of larval period (days) of *Anticarsia gemmatalis* (AG), *Chrysodeixis includens* (CI), *Helicoverpa armigera* (HA), *Spodoptera cosmioides* (SC), *S. eridania* (SE), and *S. frugiperda* (SF) larvae infesting leaf discs of IPRO and RR soybean varieties with larvae of L1 and L3 growth stages. Values with the same letter are not significantly different according to contrasts in the mixed model test (α = 0.10). Bars indicate standard error of the mean.

### Foliar area consumption

The foliar area consumed was first analyzed like the duration of larval period, considering only the larvae that reached to pupae. In the first comparison (Figure 4 Left), there was a significant effect of Stage, Species, and their interaction (p<0.0001 in all the cases). The Species effect and the interaction with Stage were due mostly to a lower consumption of L1 in *A. gemmatalis*. In the *Spodoptera* species (Figure 4 Right), there was significance of single effects Stage, Species, and Variety (p<0.0001 in the three cases), as well as the Species x Variety (p=0.0011) and Stage x Species x Variety (p=0.0039), but not from Stage x Variety (p=0.1009) nor Stage x Species (p=0.2284) interactions. The triple interaction shows that the foliar area consumed by L1 and L3 on IPRO and RR soybeans varied across species. Indeed, *S. cosmioides* had the highest consumption of all the species across larval stages and varieties, *S. eridania* consumed less foliar area overall, with a trend of higher consumption in IPRO, and *S. frugiperda* consumed the least of the *Spodoptera* spp., with the exception of the high leaf area consumed by L1 on IPRO.

**Figure 4:**
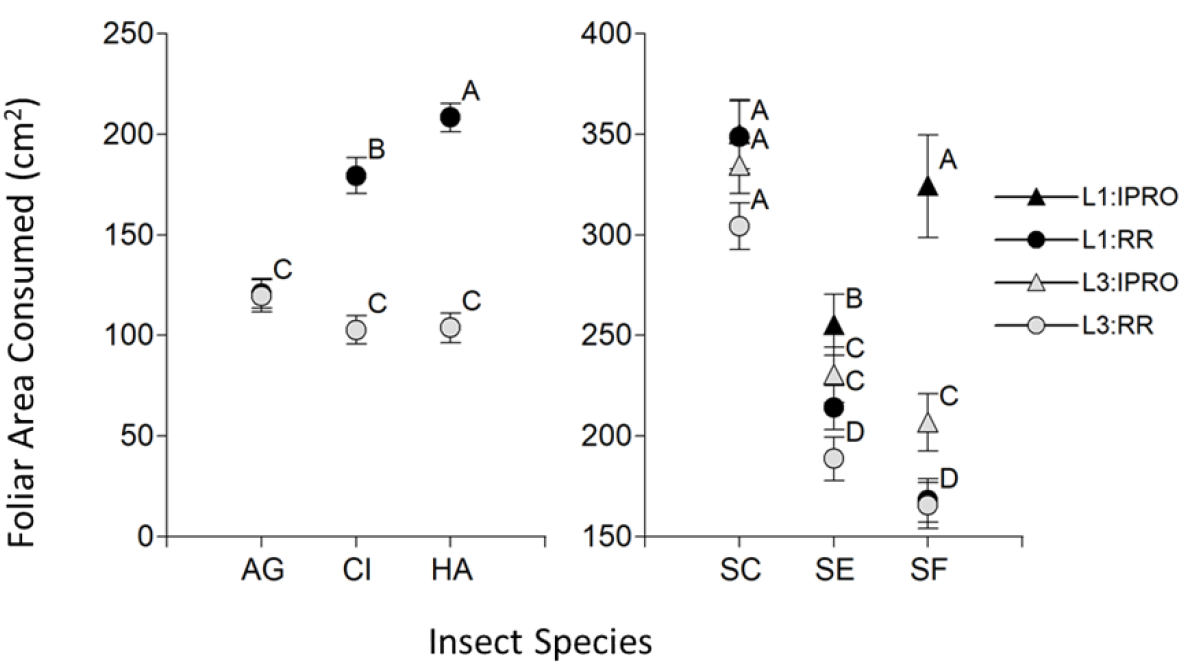
Foliar area consumed (cm^2^) of *Anticarsia gemmatalis* (AG), *Chrysodeixis includens* (CI), *Helicoverpa armigera* (HA), *Spodoptera cosmioides* (SC), *S. eridania* (SE), and *S. frugiperda* (SF) larvae (including only those that reached pupal stage) infesting leaf discs of IPRO and RR soybean varieties with larvae of L1 and L3 growth stages. Values with the same letter are not significantly different according to contrasts in the mixed model test (α = 0.10). Bars indicate standard error of the mean.

A second approach to analyze foliar area consumed included also larvae that did not reach pupal stage (Figure 5). This analysis indicated significance of Species, Variety, Stage x Species, Variety x Species (p<0.0001 in the cases), Stage x Variety (p=0.0077) and Stage x Species x Variety (p=0.0745), but not from Stage (p=0.9249). The analysis of the triple interaction showed the same pattern for the species that did not survive on IPRO variety (*A. gemmatalis, C. includens* and *H. armigera*), because larvae hardly consumed any area before being controlled. On the contrary, the pattern of foliar area consumed changed in all of the *Spodoptera* species. Indeed, in *S. cosmioides* L3 consumed more foliar area than L1, in *S. eridania* the foliar area consumed was similar in both larval sizes, and in *S. frugiperda* L1 in IPRO had the lowest consumption.

**Figure 5:**
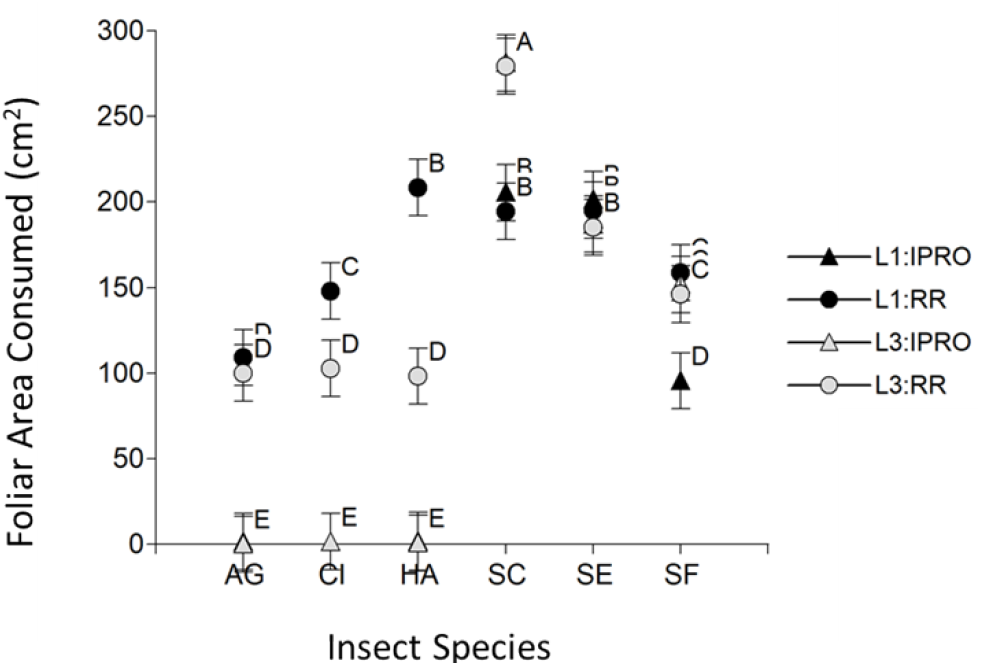
Foliar Area Consumed (cm^2^) by *Anticarsia gemmatalis* (AG), *Chrysodeixis includens* (CI), *Helicoverpa armigera* (HA), *Spodoptera cosmioides* (SC), *S. eridania* (SE), and *S. frugiperda* (SF) larvae (including both those that reached pupal stage and those that did not) infesting leaf discs of IPRO and RR soybean varieties with larvae of L1 and L3 growth stages. Values with the same letter are not significantly different according to contrasts in the mixed model test (α = 0.10). Bars indicate standard error of the mean.

The shifts in the pattern of foliar area consumption by L1 and L3 of *Spodoptera* species feeding on IPRO or RR varieties was related to the proportion of larvae reaching to pupal stage (survival). The strongest example were L1 of *S. frugiperda* on IPRO, which consumed the lowest or the highest foliar area if all the larvae or only the surviving ones were considered respectively in the analysis. This was related directly to the high mortality of L1 on IPRO (Figure 2). This same shift was also seen in *S. eridania*, in which the lower survival in IPRO compensated the higher area consumed by surviving larvae compared to RR, yielding a similar foliar area consumed in both varieties if all the larvae were considered, and also in *S. cosmioides*, comparing the consumption of L1 and L3. In this sense, Figure 6 shows that the foliar area consumed increases (5.03 cm^2^ per pupa, p<0.0001, R^2^ 0.20) with the number of pupae when all the larvae were considered (Left), but decreases if only surviving larvae (Right) are considered (−8.28 cm^2^ per pupa, p<0.0001, R^2^ 0.32).

**Figure 6:**
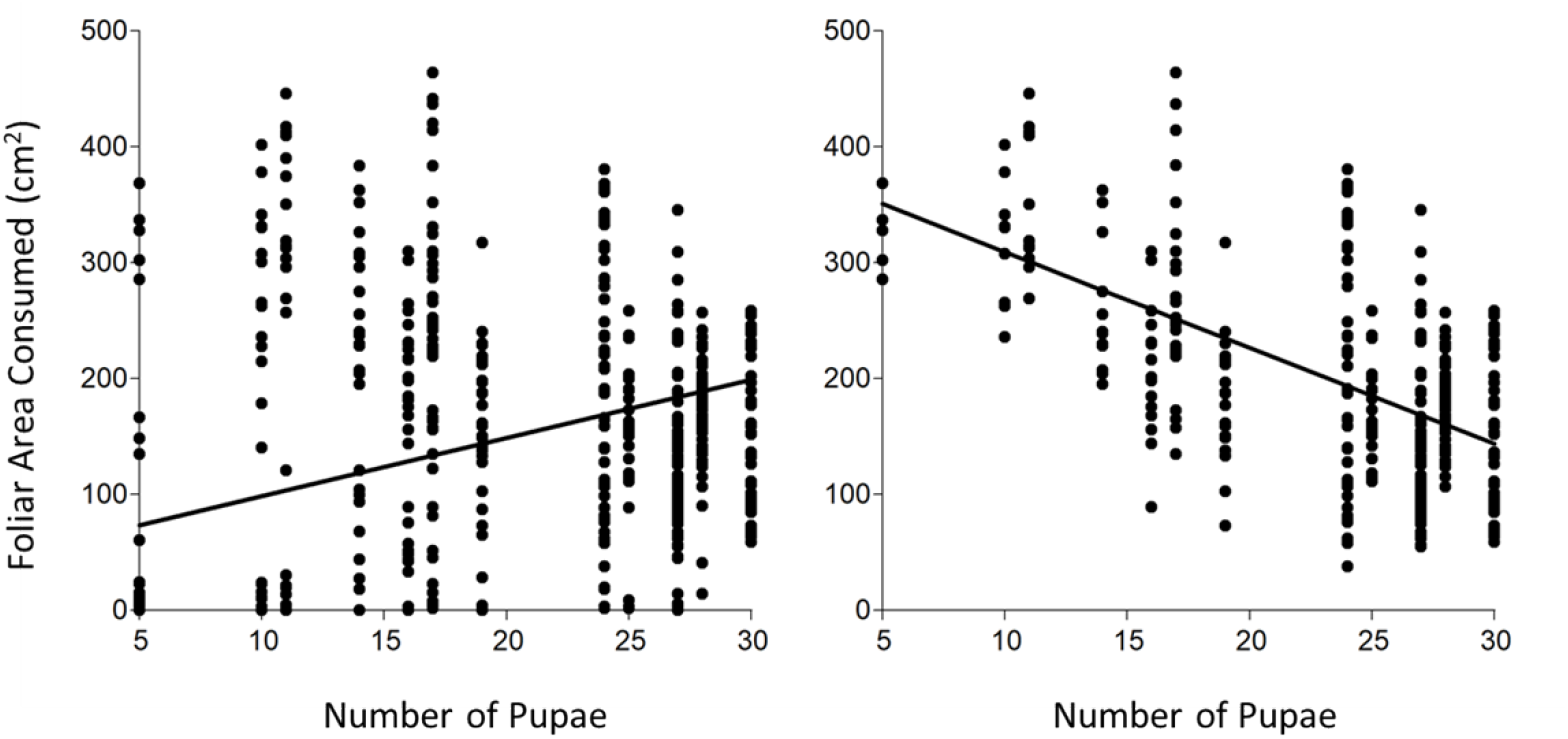
Regression of foliar area consumed (cm^2^) with number of pupae including those that reached pupal stage and those that did not (left) and only those that reached pupal stage (right). All insect species, larval stages at infestation and soybean varieties combined.

### Pupal weight

The pupal weight was analyzed like the duration of larval period. In the first comparison (Figure 7 Left), there was a significant effect of Species (p<0.0001), Stage (p=0.0003), and their interaction (p=0.0004). The Species effect was due to a larger pupal weight of *H. armigera*, and the interaction to a larger pupal weight in L3 than in L1 in *A. gemmatalis* and *C. includens* but not in *H. armigera*. The analysis of the *Spodoptera* species showed a significant effect of Species (p<0.0001), Stage (p=0.0069), and Variety (p<0.0001), and the interactions Species x Variety (p<0.0001) and Stage x Variety (p=0.0143), while the interactions Stage x Species (p=0.4932) and Stage x Species x Variety (p=0.5021) were not significant. The Species effect was because pupae of *S. cosmioides* weighted more than of the other species, and the Species x Variety interaction was due to a largest weight of pupae in RR than in IPRO in *S. cosmoides* and *S. eridania*, and to a less extent in *S. frugiperda* in both L1 (Figure 7 Right) and L3 (not shown). The Stage effect and Stage x Variety interaction were due to larger pupal weight in L3 than in L1 in *S. frugiperda* only.

**Figure 7:**
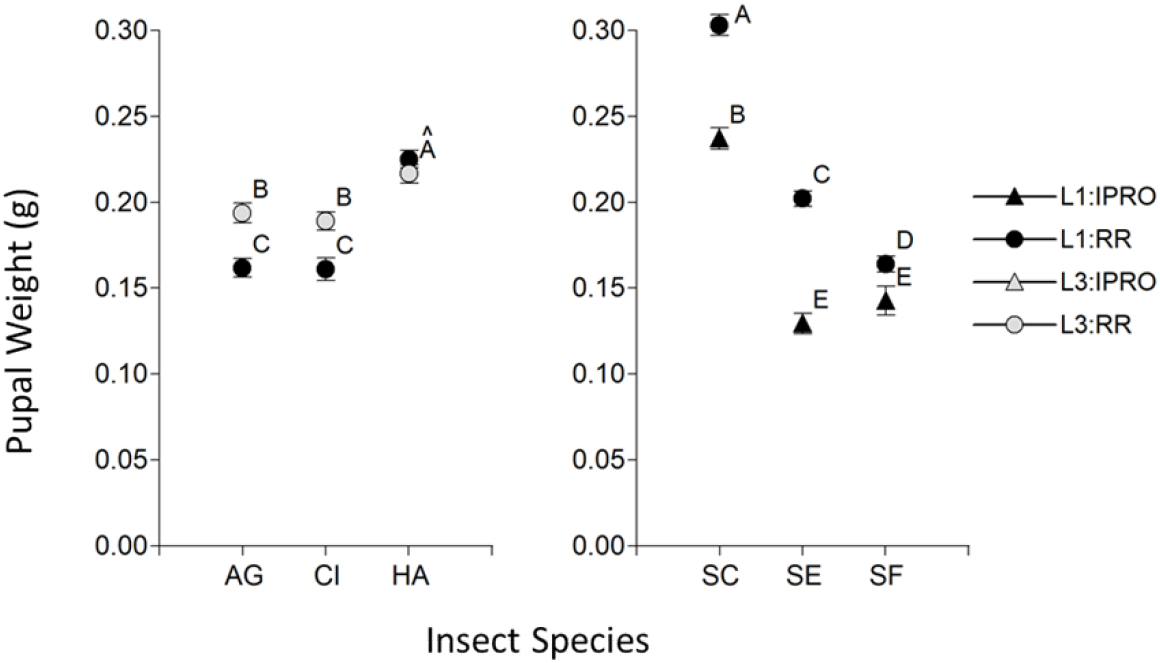
Pupal weight (g) of *Anticarsia gemmatalis* (AG), *Chrysodeixis includens* (CI), *Helicoverpa armigera* (HA), *Spodoptera cosmioides* (SC), *S. eridania* (SE), and *S. frugiperda* (SF) larvae infesting leaf discs of IPRO and RR soybean varieties with larvae of L1 and L3 growth stages. Values with the same letter are not significantly different according to contrasts in the mixed model test (α = 0.10). Bars indicate standard error of the mean.

**Figure 7:**
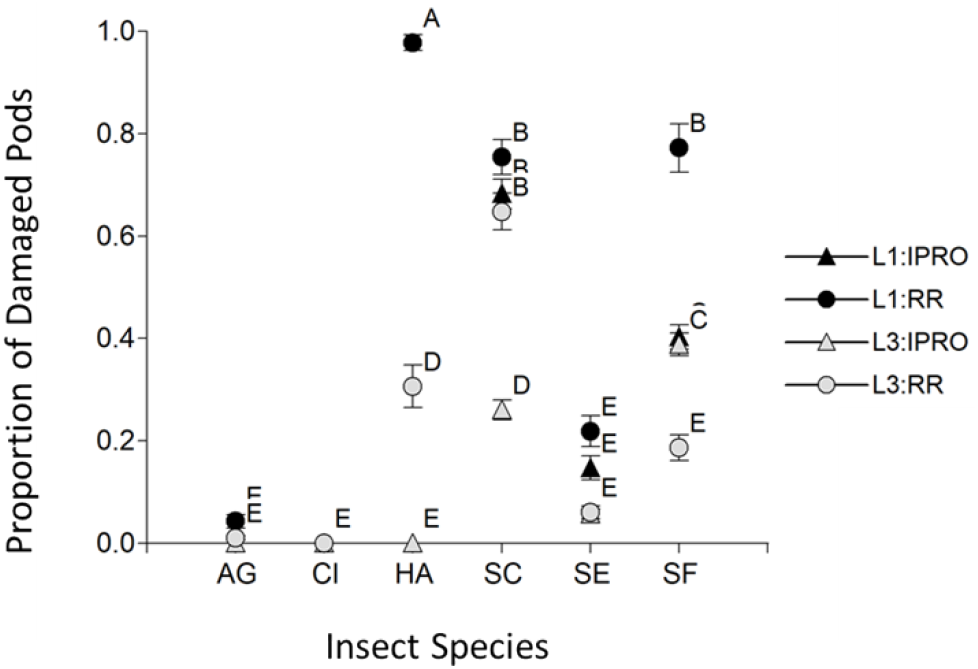
Proportion of damaged pods by *Anticarsia gemmatalis* (AG), *Chrysodeixis includens* (CI), *Helicoverpa armigera* (HA), *Spodoptera cosmioides* (SC), *S. eridania* (SE), and *S. frugiperda* (SF) larvae infesting plants at medium reproductive stages of IPRO and RR soybean varieties with larvae of L1 and L3 growth stages. Values with the same letter are not significantly different according to contrasts in the mixed model test (α = 0.10). Bars indicate standard error of the mean.

### Larval consumption at reproductive stages

The proportion of damaged pods with infestation at flowering showed no significant effect of any factor or interaction (results not shown), with less than 1% of damaged pods. On the contrary, when the infestation took place at mid grain-filling (Figure 7) there was significance of Species (p<0.0001), Stage (p=0.0395) and Variety (p=0.0008) single factors, and of Species x Variety and Species x Stage x Variety (p<0.0001 in both cases) interactions. These significances were grouped into three clusters according to the species: a first group including *A. gemmatalis, C. includens* and partially *S. eridania*, where none or very few damaged pods were found; a second group composed of *H. armigera* where pods were damaged only on RR variety, mostly in L1; and a third group consisting of *S. cosmioides* and *S. frugiperda*, where pods were damaged on both IPRO and RR varieties. These two latter *Spodoptera* species damaged more pods with infestation at L1, and L3 of *S. cosmioides* damaged more pods on IPRO, and the opposite taking place in *S. frugiperda*. The analysis of regression of grain weight with the number of damaged pods (results not shown) showed a negative relationship (0.33 g per damaged pod, p<0.0001, R^2^ 0.77).

The foliar area consumed with infestation at flowering (Figure 8 Left) showed a highly significant effect of Species, Stage, Variety, Species x Variety and Species x Stage x Variety (p<0.0001 in all the cases), as well as a Stage x Species (p=0.0019) and Stage x Variety (p=0.0042). The analysis of the triple interaction showed in that no foliar area was consumed in IPRO variety by *A. gemmatalis, C. includens* nor *H. armigera* (this latter one showing some foliar area consumed by L3 in IPRO). The *Spodoptera* species behaved differently, with *S. cosmioides* consuming less in IPRO, *S. eridania* showing no differences, and *S. frugiperda* L1 consumed more in RR and L3 in IPRO. This pattern was also seen with infestation at mid reproductive stages (p<0.0001 of all single factors and their interactions). The only major differences were the higher foliar area consumed by *S. cosmioides* in L3 in RR soybeans, a lower consumption in L1 in IPRO soybeans in *S. eridania*, and in the pattern of consumption by *S. frugiperda* (Figure 8, Right). The analysis of regression of foliar area consumed with the number of damaged pods (results not shown) showed a positive relationship (0.21 g per damaged pod, p<0.0001, R^2^ 0.26).

**Figure 8:**
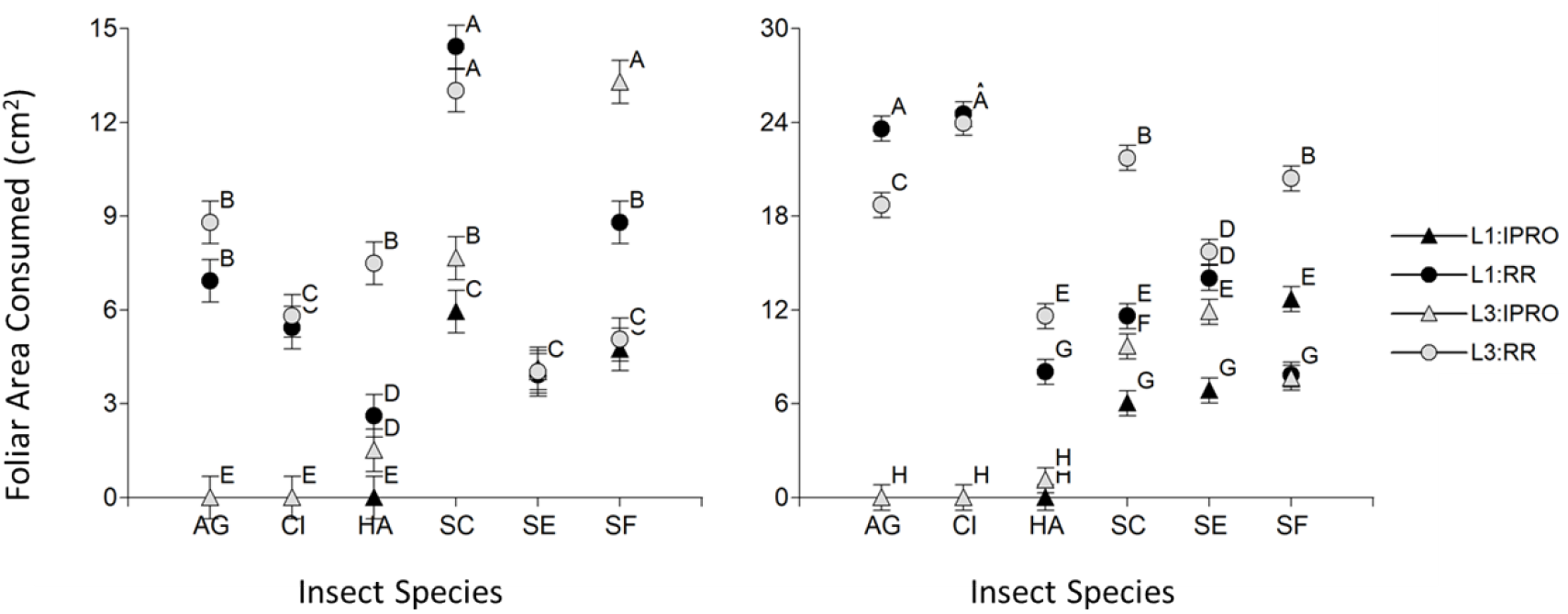
Foliar Area Consumed (cm^2^) by *Anticarsia gemmatalis* (AG), *Chrysodeixis includens* (CI), *Helicoverpa armigera* (HA), *Spodoptera cosmioides* (SC), *S. eridania* (SE), and *S. frugiperda* (SF) larvae infesting plants at early (Left) or medium (Right) reproductive stages of IPRO and RR soybean varieties with larvae of L1 and L3 growth stages. Values with the same letter are not significantly different according to contrasts in the mixed model test (α = 0.10). Bars indicate standard error of the mean.

## Discussion

Survival of larvae was high in all the species tested, similar to previous reports in *C. includens* (Kidd and Orr 2001, Barrionuevo et al. 2012, Specht et al. 2019), *H. armigera* (Gomes et al. 2017), *S. eridania* (Santos et al. 2005, Ramirez and Gómez 2010, Favetti et al. 2015), *S. frugiperda* (Silva et al. 2017a, Montezano et al. 2019). The lower survival of *S. cosmioides* was seen before (Bavaresco et al. 2003, Silva et al. 2017b), although some findings in artificial diet (Specht and Specht 2016) indicated higher survival. The survival of larvae up to pupal stage was affected in three ways: larval size at infestation, soybean variety, and their interaction. Changes in survival due to larval size were seen as higher survival by L3 than L1 in *C. includens* and *S. cosmioides*. This could be related to differences in the dynamics of survival of these species, or due to an effect of the change from feeding in artificial diet (where they fed until the infestation) to the foliar discs as found before (Reid and Greene 1973). The changes in survival attribuited to soybean variety were related to the high efficacy of the IPRO biotechnology to control *A. gemmatalis, C. includens* and *H. armigera* larvae (MacRae et al. 2005, McPherson and MacRae 2009, Yu et al. 2013, Azambuja et al. 2015), while those of *Spodoptera* spp. were controlled partially (*S. eridania* and *S. frugiperda*) or not (*S. cosmioides*), as it was formerly reported (Barros et al. 2010, Bortolotto et al. 2014, Bernardi et al. 2014, Silva et al. 2016, Murúa et al. 2018). Lastly, the interaction of larval size and soybean variety, changed the survival of *S. frugiperda*, expressed as a lower survival of L1 in IPRO than in RR variety, while no differences were seen in L3. This finding has not been documented yet to our knowledge, and may deserve further consideration in the future, since survival of larvae plays a role in foliar area consumed, and hence to estimate EIL (Hutchins et al. 1988).

Length of larval period was similar to previous reports run at similar temperatures (Kidd and Orr 2001, Bavaresco et al. 2003, Barrionuevo et al. 2012, Silva et al. 2016, Specht and Specht 2016). Besides the obvious effect of larval size at the time of infestation, there were differences among the species tested, having *A. gemmatalis* the shortest, and the *Spodoptera* species the longest duration.

Larval period in *Spodoptera* species lasted longer in IPRO variety, mainly in small larvae of *S. eridania* and *S. frugiperda* as reported before (Bortolotto et al. 2014), while *S. cosmioides* was less affected, in agreement with previous work on this species (Silva et al. 2016). This finding may contribute to a better decision-making to control foliar pests in IPRO varieties, because a slower larval growth provides more time to decide the need of control with insecticides.

The range of foliar area consumed found in this work is similar to earlier reports in *A. gemmatalis* (Bueno et al. 2011), *C. includens* (Reid and Greene 1973), *S. cosmioides* (Bueno et al. 2011), *S. eridania* (Santos et al. 2005) and *S. frugiperda* (Bueno et al. 2011) including larvae that pupated and those that did not. These results confirm that *S. cosmioides* consumes about twice foliar area than the other species, and so has to be considered separately to determine economic thresholds. Soybean variety also played a major role on foliar area consumption, since IPRO variety reduced almost entirely this consumption by *A. gemmatalis, C. includens* and *H. armigera*; but only in a small amount in the *Spoodoptera* species as reported before (Bortolotto et al. 2014, Silva et al. 2016, Murúa et al. 2018).

Pupal weight was similar to previous results in the species studied (Kidd and Orr 2001, Santos et al. 2005, Bernardi et al. 2014, Montezano et al. 2014, Silva et al. 2017a). Pupae of *Spodoptera* spp. on IPRO weighted less than in RR variety, with a smaller difference in *S. cosmioides*, contradicting previous reports that showed similar pupal weight in both varieties by these *Spodoptera* species (Bernardi et al. 2014, Silva et al. 2016). The lower pupal weight found in this study was related to a longer larval period in IPRO, althought the foliar area consumed was similar in both varieties. For this reason, we consider that this matter deserves a deeper analysis in the future, since it would help to better understand the fitness of these species on IPRO soybeans.

In our work, when infestation took place at flowering (R1-R2), there were no damaged pods with the insect pressure used (2 larvae per plant). This could be due to the large size of the plants and the high number of flowers at the time of infestation compared to the insect pressure, considering also that most flowers naturally abort in the absence of insect damage. In turn, the proportion of damaged pods during mid-grain filling varied primarily with the feeding habit of each species. On one hand, there were no (*A. gemmatalis* and *C. includens*) or few (*S. eridania*) damaged pods in species not considered consistently as pod feeders. One aspect to consider is that no damage of reproductive structures by *A. gemmatalis* was seen in this work. This agrees with the most usual behavior of this species, although some other reports (Gamundi 1988) showed that this species feeds from soybean pods at R3-R4 stages. Further research on this area may be useful, due to the high prevalence of this species in South America. On the other hand, the species *H. armigera, S. cosmioides* and *S. frugiperda* did damaged pods, agreeing with previous results (Adams et al. 2016) of their feeding behavior. The damage to pods resembled the one to foliar area in *H. armigera* damaging RR variety only, and *S. cosmioides* and *S. frugiperda* damaging both varieties. The results in *H. armigera* confirm previous work (MacRae et al. 2005), establishing that species controlled by IPRO during vegetative stages do not damage reproductive structures in the larval sizes and insect pressure used in this study. In turn, while no major differences were seen in the damage to pods by *S. cosmioides* in larval sizes and varieties, in *S. frugiperda* damage to pods was higher in RR than in IPRO in both larval sizes. These results resemble those of foliar area consumed, and agree with previous work indicating higher tolerance of *S. cosmioides* than *S. frugiperda* (Bernardi et al. 2014, Silva et al. 2016, Murúa et al. 2018) to IPRO varieties.

An important aspect to determine the economic injury level (EIL) is the consumption of vegetative and reproductive structures by species able to feed from both of them. In this work, the presence of reproductive structures did not consistently change the foliar area consumption across species, larval sizes and varieties. For instance, *S. cosmioides* was the species with the highest foliar area consumption during vegetative stages, but during reproductive stages this species reduced the consumption of foliar area in favor to feeding from pods. In turn, in *H. armigera* and *S. frugiperda* this shift was seen only on L1 of in RR variety. In this sense, future areas of research could focus in understanding the nature of this difference across species.

Integrating the consumption of foliar area adjusted by survival and proportion of damaged pods to obtain an EIL for each species, we identified the following behaviors: a) *A. gemmatalis* and *C. includens*, causing only defoliation, limited to the RR variety, having the lowest consumption of foliar area; b) *S. eridania*, causing majorly defoliation in both RR and IPRO varieties, consuming twice as much foliar area than the abovementioned species; c) *H. armigera*, limited to the RR variety causing an intermediate consumption of foliar area and the highest damage to pods; d) *S. cosmioides* and *S. frugiperda*, causing the highest (*S. cosmioides*) or intermediate (*S. frugiperda*) damage in both defoliation and to pods in both varieties. Despite having a similar range of feeding, these two latter species differed quantitatively in the ability to feed from IPRO variety, with *S. cosmioides* causing similar damage to both varieties and *S. frugiperda* showing a trend of lower damage in IPRO. This clasification contributes to a sustainable recommendation of insecticide use, taking into account the behavior of these species that are major soybeans pests in South America.

